## Supplement Chen at al for "Engineering protein-specific proteases: targeting active RAS"

### Supplemental Results

**RAS cleavage in human cell culture** To date we have only tested one RAS protease in human cells, but we have established that a RAS-specific protease can self-activate in human cells, locate KRAS at the plasma membrane, and cleave it as indicated by the presence of the eGFP fusion product and the precipitous disappearance of KRAS in **Fig. 8**. Moreover, as observed in the *E. coli* experiments, cleavage of KRAS at switch 2 in mammalian cells creates a highly unstable product as evidenced by the disappearance of eGFP (**Fig. 8)**. Although RAS isoforms are highly abundant in human cells (~300,000 RAS molecules (150nM) in colorectal cancer cells) (1), the proteases developed in *E. coli* will need to be adapted to the new environment if precise control of RAS signaling is the goal. Perhaps the more important point, however, is that the general principles learned from *E. coli* makes re-programming cultured human cells appear feasible. We have also used a targeted nanocarrier system to deliver protease to the cytoplasm of cells (Supplement Protein Delivery) as a potential means to control the amount of active protease in cells and define the amount required to eliminate a particular RAS isoform or mutant.

**Supplemental Materials and Methods**

**Supplement NMR spectroscopy** Samples for three-dimensional experiments were approximately 0.25 mM RAS in 20 mM HEPES, 50 mM NaCl, 5 mM $MgCl_2$, and 1 mM TCEP, pH 7.4. All assignment experiments were collected at 25°C. The following standard three-dimensional heteronuclear experiments were acquired: HNCACB, CBCA(CO)NH, HN(CA)CO, HNCO. Triple resonance experiments were performed with 25% non-uniform sampling (2). Amide chemical shift perturbations between the GDP and GMPPNP-bound forms of G12V-HRAS, $\Delta\delta_{total}$, were calculated from the equation $\Delta\delta_{total} = [(W_H\Delta\delta_H)_2 + (W_N\Delta\delta_N)_2]^{1/2}$, where $\Delta\delta_H$ and $\Delta\delta_N$ are the proton and nitrogen shift differences for a given resonance, respectively, and $W_H$=1 and $W_N$=0.2 are weighting factors. Two-dimensional heteronuclear $\{^1H\}$-$^{15}N$ steady state NOE experiments were collected with a gradient-selected, sensitivity-enhanced pulse sequence

(3). Spectra were recorded with and without saturation in an interleaved manner employing a 5 s recycle delay. The heteronuclear NOEs were calculated from the NOE_on/NOE_off ratio and standard deviations were obtained from measured background noise levels. $^{15}N$ $R_1$ and $R_2$ measurements were carried out using gradient-selected, sensitivity enhanced 2D $^{1}H$-$^{15}N$ HSQC experiments (3) with water flip-back modifications for solvent suppression. $^{15}N$ $R_1$ experiments were acquired with variable delay times of 10, 150, 300, 400, 600, 900, and 1200 ms. $^{15}N$ $R_2$ experiments were acquired with delay times of 8, 16, 24, 32, 40, 48, 64, 80, 96, and 130 ms. $R_1$ and $R_2$ values were obtained from single exponential decay fitting with error estimates for $R$ using Sparky. Generalized order parameters ($S^2$) were extracted from the $^{15}N$ relaxation data utilizing the Modelfree program (4). Spectra were processed using NMRPipe (5) and analyzed with Sparky (6).

**Supplemental crystallization, data collection and processing** For the RASProtease(I)-YSAM complex, RASProtease(I) crystals were soaked overnight in mother liquor supplemented with the YSAM peptide at a final concentration of 2.1 mM. RASProtease(I) crystallizes in a variety of conditions and for the YSAM complex, the mother liquor for the YSAM complex contained 0.1 M TRIS-HCl pH 8.0, 20% PEG 6K, 0.2 M NaCl. Crystals were transferred to the crystallization condition supplemented with 15% glycerol and 2.1 mM YSAM peptide and flash frozen in liquid nitrogen prior to data collection. Native data up to 1.2 Å resolution were collected at Stanford Synchrotron Radiation Laboratory (SSRL) beamline 12-2. The data were reduced using mosflm (7) and Aimless (8) from the CCP4 program suite (8). A similar procedure was used to generate the RASProtease(I) -QEEYSAM complex. The mother liquor for these crystals was 0.1 M HEPES pH 7.0, 20% PEG 6K, 0.2 M NaCl. Data for the RASProtease(I) -QEEYSAM complex were collected to 1.63 Å using in-house X-ray diffraction resources. A summary of the data collection statistics is provided in **Table S3**.

Similarly, purified Protease1(N) was concentrated to 12 mg/mL in a buffer composed of 5 mM HEPES pH 7.0, 10 mM Azide, and 0.53 mM LFRAL pentapeptide. Initial crystallization screening at 17 °C using in-house resources identified a number of potential conditions, the best of which was 30% PEG 8K, 0.2M NaCl, and 0.1 M Imidazole pH 8.0. This condition was subsequently refined to 0.1 M Imidazole pH 8.4, 0.2 M NaCl, and 25% PEG 8K. Like the RASProtease(I) crystals, these crystals belong to space group $P4_12_12$ and have unit cell dimensions a=b=58.7 Å, c=125.7 Å, $\alpha=\beta=\gamma=90°$. A crystal from this condition was cryo-protected using mother liquor mixed with glycerol to give a final glycerol concentration of 17%, cryocooled directly in the gaseous nitrogen stream of the X-ray source, and diffraction data were collected up to 1.6 Å resolution. The data were reduced using the D*Trek package (9).

**Supplemental structure determination, model building and refinement** Initial phases for RASProtease(I) and the RASProtease(I) -YSAM complex were determined by molecular replacement using the program Molrep (10) using the coordinates of the subtilisin chain in a subtilisin-prosegment complex structure (PDB ID: 1SPB). Prior to molecular replacement, all solvent molecules and ions were removed from the search model. The structure was refined using Refmac5 (11-15) from CCP4 program suite. Iterative cycles of model building using COOT (16-19) yielded structures with $R_{work}/R_{free}$ of 0.14/0.18 for RASProtease(I) , 0.09/0.11, for the RASProtease(I) -YSAM complex, and 0.15/0.18 for the RASProtease(I) -QEEYSAM complex. A summary of the refinement statistics for all structures is provided in **Table S3**. The refined structures were deposited in the PDB (Accession Codes: 6U9L, 6UAI, 6UAO).

Initial phases for the Protease1(N) -LFRAL complex were determined by using the program Molrep (10) using the coordinates of a previously-determined subtilisin crystal structure (PDB ID: 3F49). Prior to molecular replacement, all solvent molecules and ions with B factors greater than 20 $Å^2$ were removed from the search model. The structure was refined using Refmac5 (11-15) from the CCP4 program suite. Iterative cycles of model building using COOT (16-19) yielded a structure with $R_{work}/R_{free}$ of 0.16/0.18. A summary of the refinement statistics for the structure is provided in **Table S3**. The refined structure was deposited in the PDB (Accession Code: 6UBE).

**Analysis of KRAS cleavage in cells** To ensure a relatively uniform amount of nitrite in cells transfected with the designed protease, 1 mM sodium nitrite was added to the media along with doxycycline. Following this incubation, the media in each well was replaced with PBS and the cells from each well were harvested. eGFP-KRAS and/or its cleavage products were isolated from the cells using a GFP-trap_A immunoprecipitation kit (Chromotek). Briefly, cells were lysed in 10 mM TRIS-HCl pH 7.5, 150 mM NaCl, 0.5 mM EDTA, 1% (v/v) Triton X-100 on ice according to the manufacturer's recommendations. After centrifugation to remove insoluble cell debris, the lysates were incubated with anti-GFP beads that had been equilibrated in 10 mM TRIS-HCl pH 7.5, 150 mM NaCl, 0.5 mM EDTA overnight with gentle agitation at 4 °C. The beads were then washed three times with 10 mM TRIS-HCl pH 7.5, 150 mM NaCl, 0.5 mM EDTA. Following removal of the wash buffer, the beads were resuspended in SDS-PAGE running buffer and boiled for 10 minutes at 95 °C prior to analysis by Western blotting. The same lysates were then used for anti-FLAG pulldowns using a FLAG immunoprecipitation kit (Sigma) according to the manufacturer's instructions. The procedure is essentially as described above, except that the wash buffer is 50 mM TRIS-HCl pH 7.5, 150 mM NaCl and the samples were boiled for three minutes at 95 °C prior to analysis by Western blotting.

**Microscopy** Six-well tissue culture plates (Thermo Fisher Scientific cat. No. 140675) were imaged with a Zeiss LSM 710 AxioObserver microscope with plate adapter (cat. no. 451353-0000-000). Eight bit images were taken with an EC Plan-NeoFluar 10x/0.30 M27 objective. A 16% power, 514nm wavelength was used to excite GFP conjugated to protein. Four images from each well near the center were taken to minimize reflection and light artifacts at 24, 48 and 72 hours post transfection resulting in 24 images at each timepoint.

**Quantifying Fluorescent Cells** A representative timepoint image was opened in Zen Lite (Blue Edition v. 2.6, Zeiss). The green channel histogram was set from default 255 to 50. This increased the brightness before being exported into .tiff format. Images in the timepoint folder were processed similarly using "Batch" mode. Representative images were opened in Photoshop (CC v. 20.0.0, Adobe). Each was converted to black and white. Green channel was set to 300%. Next the images were flattened and inverted. Exported images were grayscale, 8-bit, .tiff files after using "Actions" (batch processing) in Photoshop. Images were imported into FIJI[1] (Image J, NIH) and counted with the "Analyze Particles" module. Size was set to 0.01-0.04 with circularity equal to 0.00-0.50. The settings were saved to a "Macro" and used as a batch process on all images in the folder. Prism (v8.0, GraphPad) was used to plot the counts. Additional information can be accessed at: https://www.protocols.io/view/quantifying-fluorescent-cells-in-mammalian-cell-ti-y2nfyde .

**Definition of mutants** *Subtilisin:* SBT189 (3BGO.pdb) is our starting subtilisin whose engineering has been described earlier. SBT189 denotes subtilisin from *Bacillus amyloliquefaciens* with the following mutations: Q2K, S3C, P5S, S9A, I31L, D32A, K43N, M50F, A73L, Δ75-83, Y104A, G128S, E156S, G166S, G169A, S188P, Q206C, N212G, Y217L, N218S, T254A, Q271E (*21, 23-26*).

**Measurement of activity of protease** Synthetic peptide-AMC (7-amino-4-methylcoumarin (AMC, Ex: 350 nm, Em: 450 nm) was purchased from AnaSpec Inc. Concentrations of the AMC substrates were determined by absorbency at 324 nm using an extinction coefficient of 16 $mM^{-1}cm^{-1}$. Reaction kinetics of AMC substrates were measured using a KinTek Stopped-Flow Model SF2001 (Ex: 380 nm, Em: 400 nm cutoff filter). Kinetic data were fit using KinTek Global Explorer software obtained from the KinTek Corporation website (www.kintek-corp.com).

**Table S1: Data Collection and Refinement Statistics**

| | RASProtease(I) | RASProtease(I) **+ YSAM** | Protease1(N) **+ LFRAL** | RASProtease(I) **+ QEEYSAM** |
|---|---|---|---|---|
| PDB code | 6U9L | 6UAI | 6UBE | 6UAO |
| Wavelength (Å) | 1.5418 | 0.97946 | 1.5418 | 1.5418 |
| Space Group | $P4_12_12$ | $P4_12_12$ | $P4_12_12$ | $P4_12_12$ |
| Unit Cell (Å) | a=58.65 b= 58.65 c=124.75 | a=58.64 b= 58.64 c=125.17 | a=58.65 b= 58.65 c=125.67 | a=58.51 b= 58.51 c=125.09 |
| Resolution (Å) | 53.3 - 1.7 | 53.1 – 1.2 | 19.3 – 1.6 | 42.7 – 1.63 |
| Unique Reflections | 24845 | 69948 | 29454 | 28012 |
| Completeness | 100.0 (100.0) | 100.0 (100.0) | 98.9 (91.3) | 100 (99.8) |
| Multiplicity | 27.2 (26.6) | 22.6 (22.6) | 5.8 (3.2) | 13.5 (13.2) |
| $\langle I/\sigma(I)\rangle$ | 26.7 (8.3) | 30.6 (25.4) | 25.2 (7.6) | 20.6 (2.8) |
| $R_{meas}$ | 0.099 (0.424) | 0.105 (0.120) | 0.041 (0.158) | 0.093 (0.808) |
| $CC_{1/2}$ | 0.999 (0.980) | 0.997 (0.996) | n/a | 0.999 (0.872) |
| Wilson B factor (Å$^2$) | 13.6 | 5.3 | 13.8 | 13.2 |
| **Refinement** | | | | |
| $R_{work}$ | 14.7 | 10.9 | 12.8 | 14.9 |
| $R_{free}$ | 17.4 | 12.5 | 15.8 | 18.0 |
| No. of non-H atoms | | | | |
| Protein | 1854 | 1976 | 1943 | 1940 |
| Waters | 185 | 321 | 299 | 242 |
| Potassium | 3 | | | |
| Sodium | | 3 | 2 | 1 |
| Chloride | | 1 | | 1 |
| Ethylene glycol | | 20 | | 20 |
| Glycerol | 12 | | 18 | |
| Polyethylene glycol | | 7 | | |
| Thiocyanate | 15 | | | |
| Azide | | | 12 | |
| Average B factor (Å$^2$) | 14.4 | 9.6 | 15.3 | 15.4 |
| *R.m.s. deviations* | | | | |
| Bonds (Å) | 0.02 | 0.02 | 0.02 | 0.01 |
| Angles (°) | 2.1 | 2.2 | 2.0 | 1.8 |
| Ramachandran favored (%) | 98 | 96 | 98 | 97 |
| Ramachandran outliers (%) | 0.4 | 1 | 0.4 | 0.4 |

**Supplemental Figures**

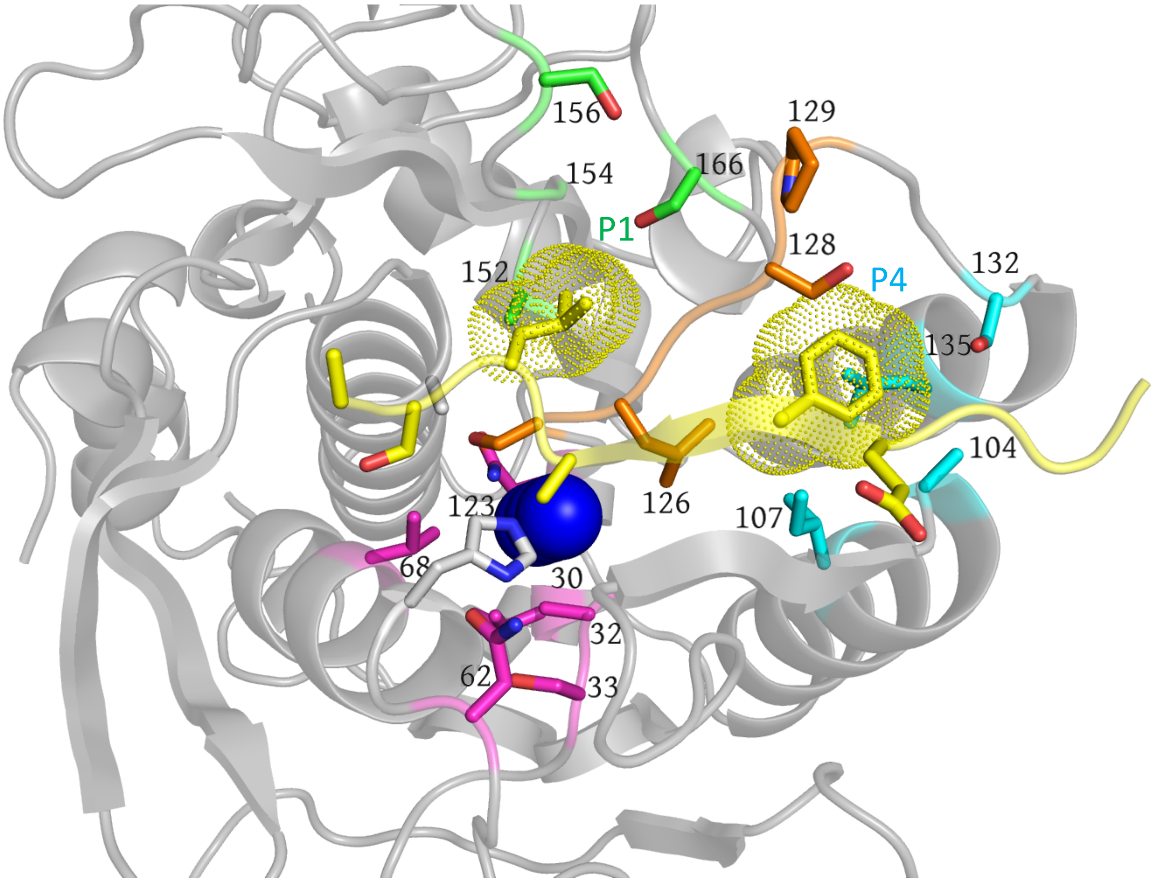


**Fig. S1:** Sites modeled for mutagenesis in the P1 pocket are in green, sites in the P4 pocket in cyan, and sites in the anion pocket are in violet. P1 leucine and P4 phenylalanine are shown with dot surfaces. The three binding sites are interconnected by common amino acids in the region from 123-129. These amino acids are in orange. Most subtilisin contacts are with the first five substrate amino acids on the acyl side of the scissile bond (denoted P1 through P5, numbering from the scissile bond toward the N-terminus of the substrate and the first amino acid on the leaving group side. Model based on 3BGO.pdb (*25*).

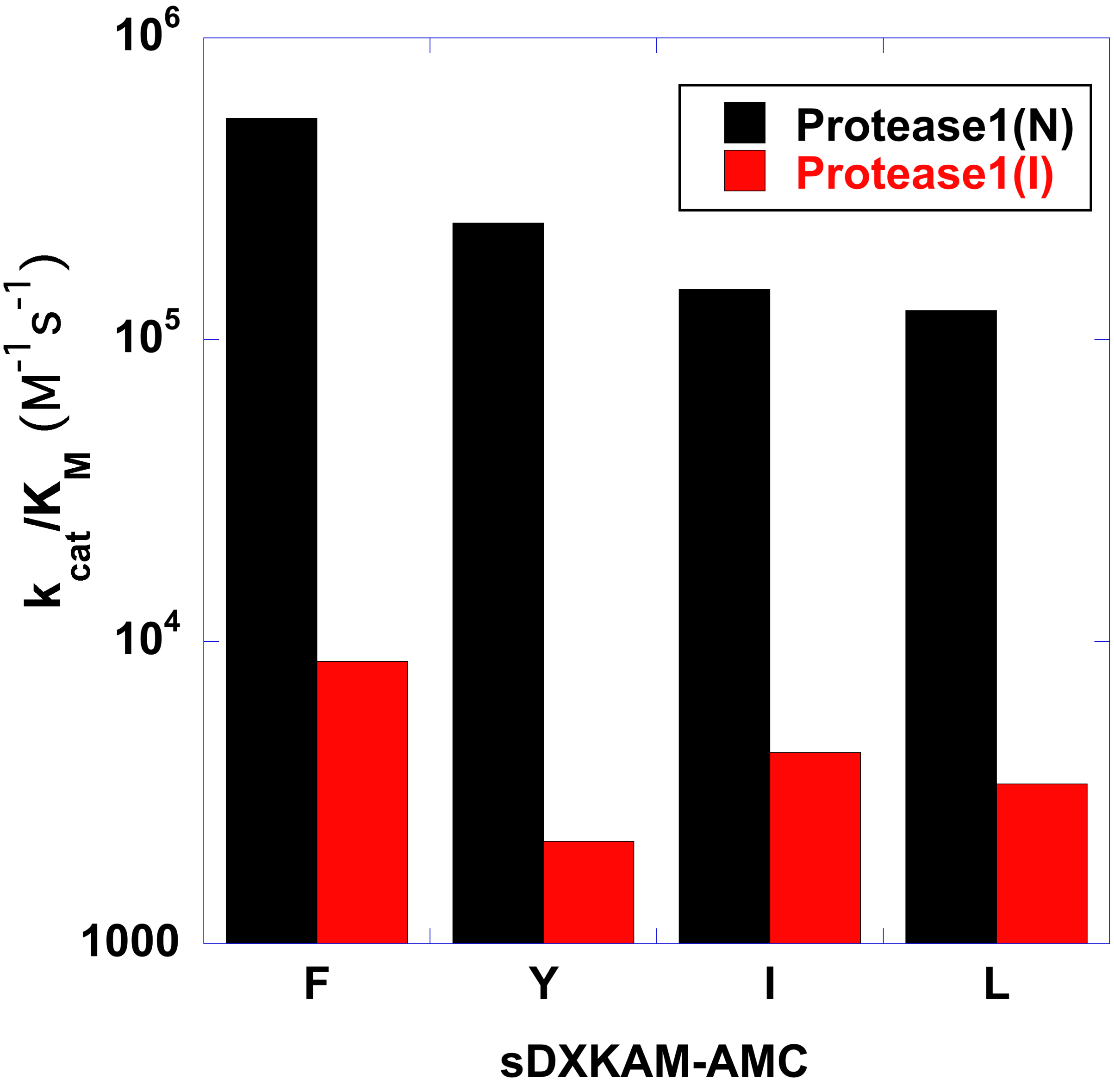


**Fig. S2:** First generation proteases evaluated with DXKAM-AMC substrate series. $k_{cat}/K_M$ values are shown for hydrophobic amino acids at P4. Protease1(N): 10mM nitrite; Protease1(I): 10mM imidazole.

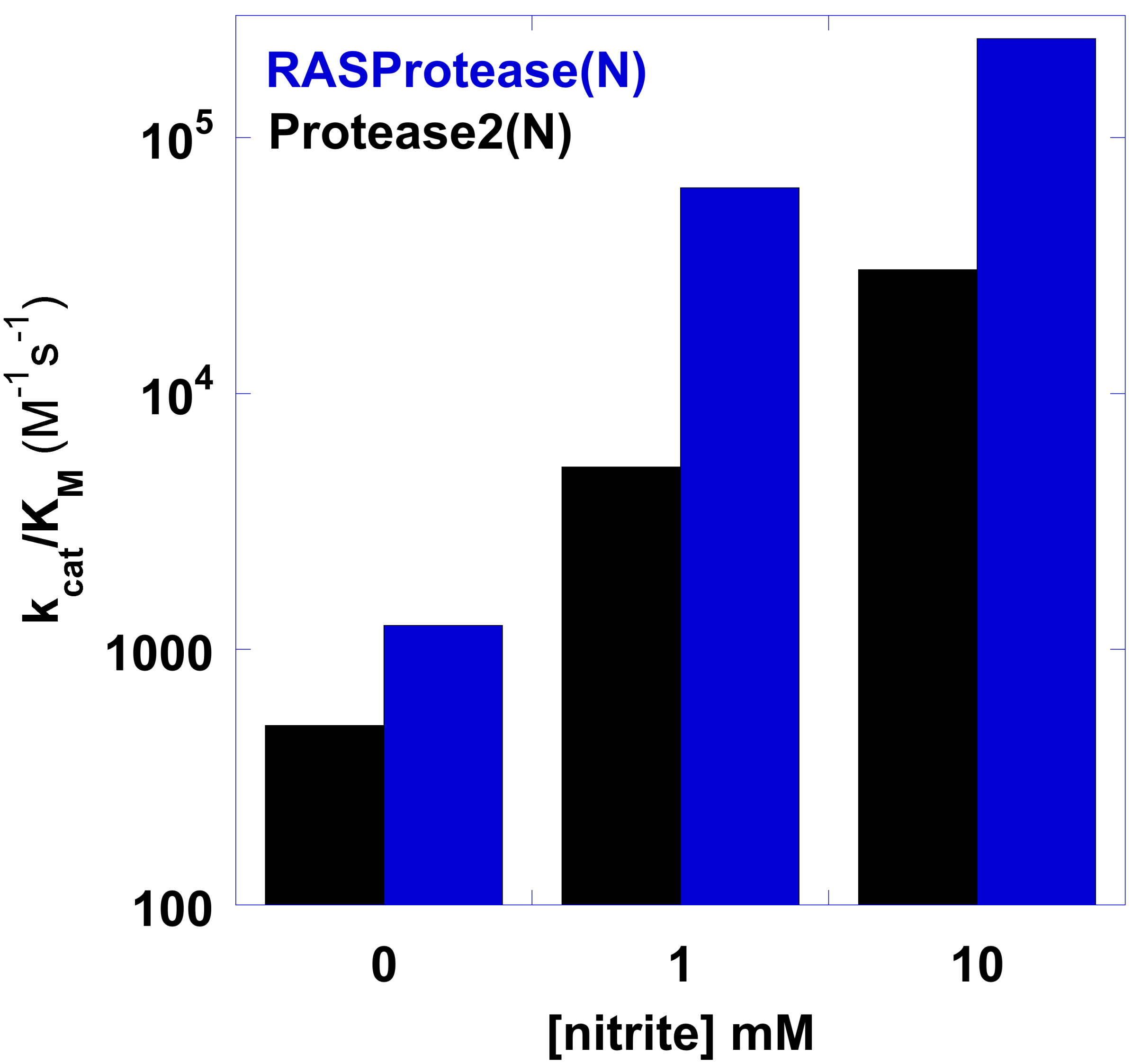


**Fig. S3:** $k_{cat}/K_M$ as a function of nitrite concentration for Protease2(N) and RASProtease(N) with the substrate QEEYSAM-AMC.

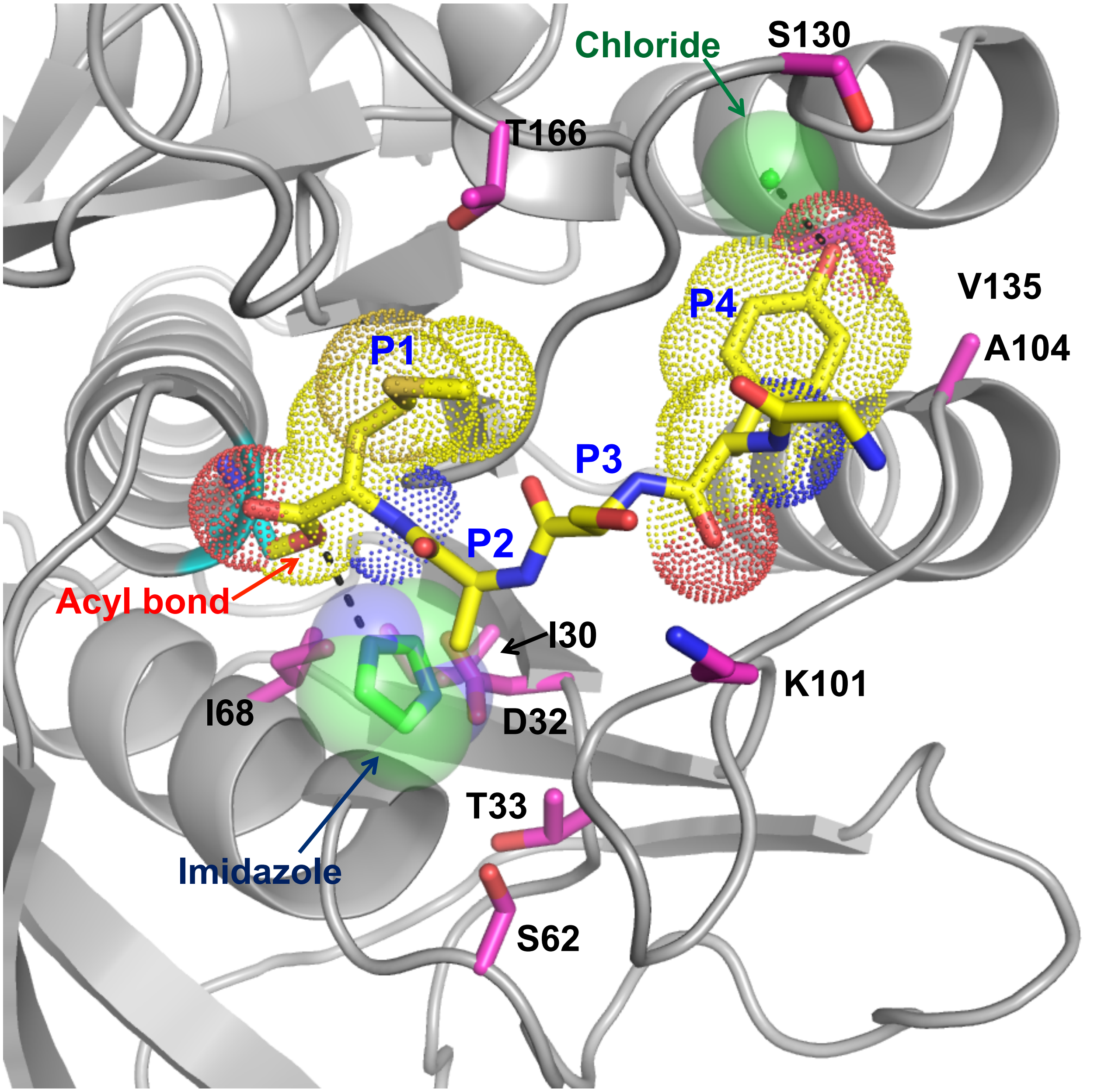


**Fig S4:** RASProtease(I) QEEYSAM with imidazole (green) modeled in place of three waters. New amino acids from the design process are in magenta. Substrate is in yellow. Only P1 to P4 are shown for clarity. An ion (modeled as chloride, pale green sphere) interacts with the hydroxyl groups of the P4 Tyr and Y171 of RASProtease(I).

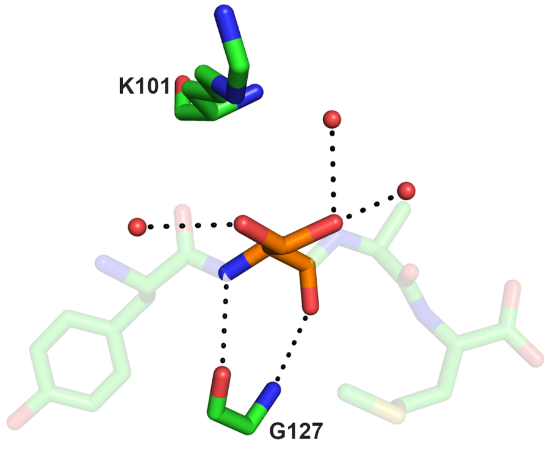


**Fig. S5:** Changes to residue 101 affect substrate preference at P3 for RASProtease(I) (left, P3=S) versus Protease1(N) (right, P3=R). Only P3 for the bound peptide is shown as solid sticks (orange) for clarity. The remaining residues are shown as semitransparent sticks.

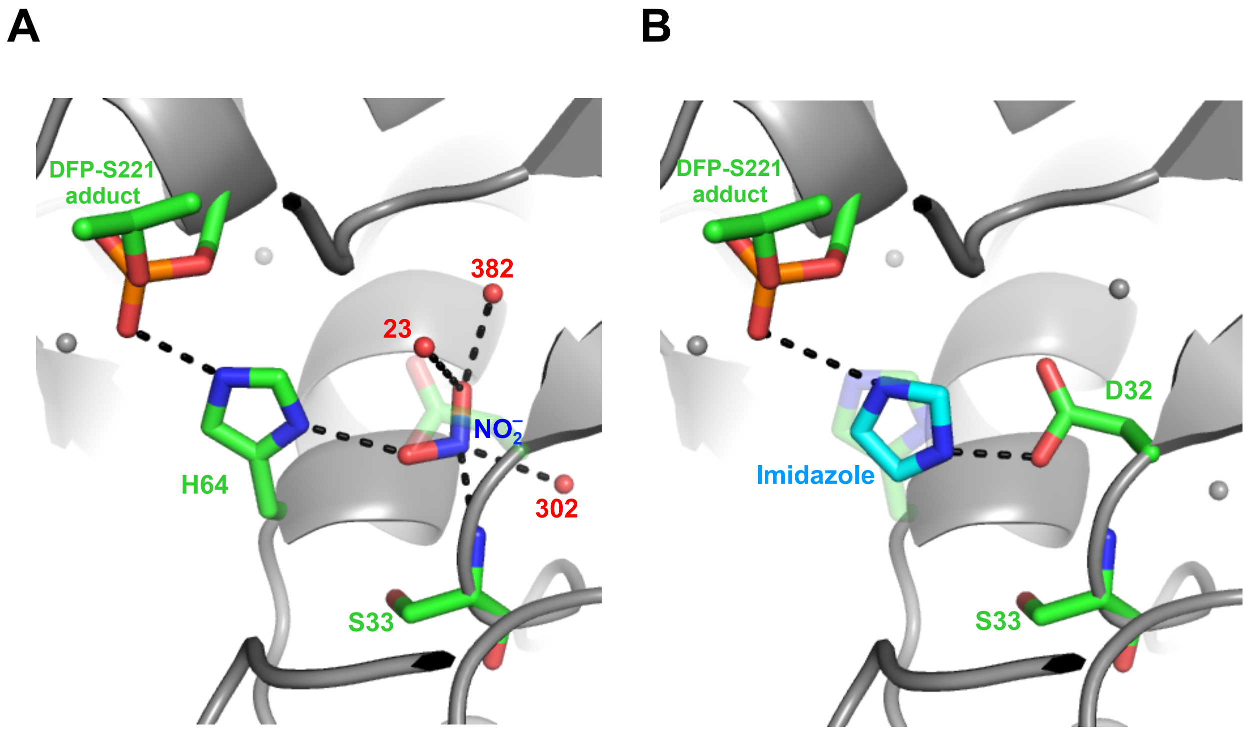


**Fig. S6:** Catalytic triad with overlaid co-factor shows how cofactors substitute for the mutated catalytic residue: A – nitrite (modeled based on 1SUE.pdb), B – imidazole (from YRGLIM structure – Bryan, Orban, and Toth, unpublished). Water molecules coordinating to nitrite in A are shown as red spheres. DFP = diisopropyl fluorophosphate.

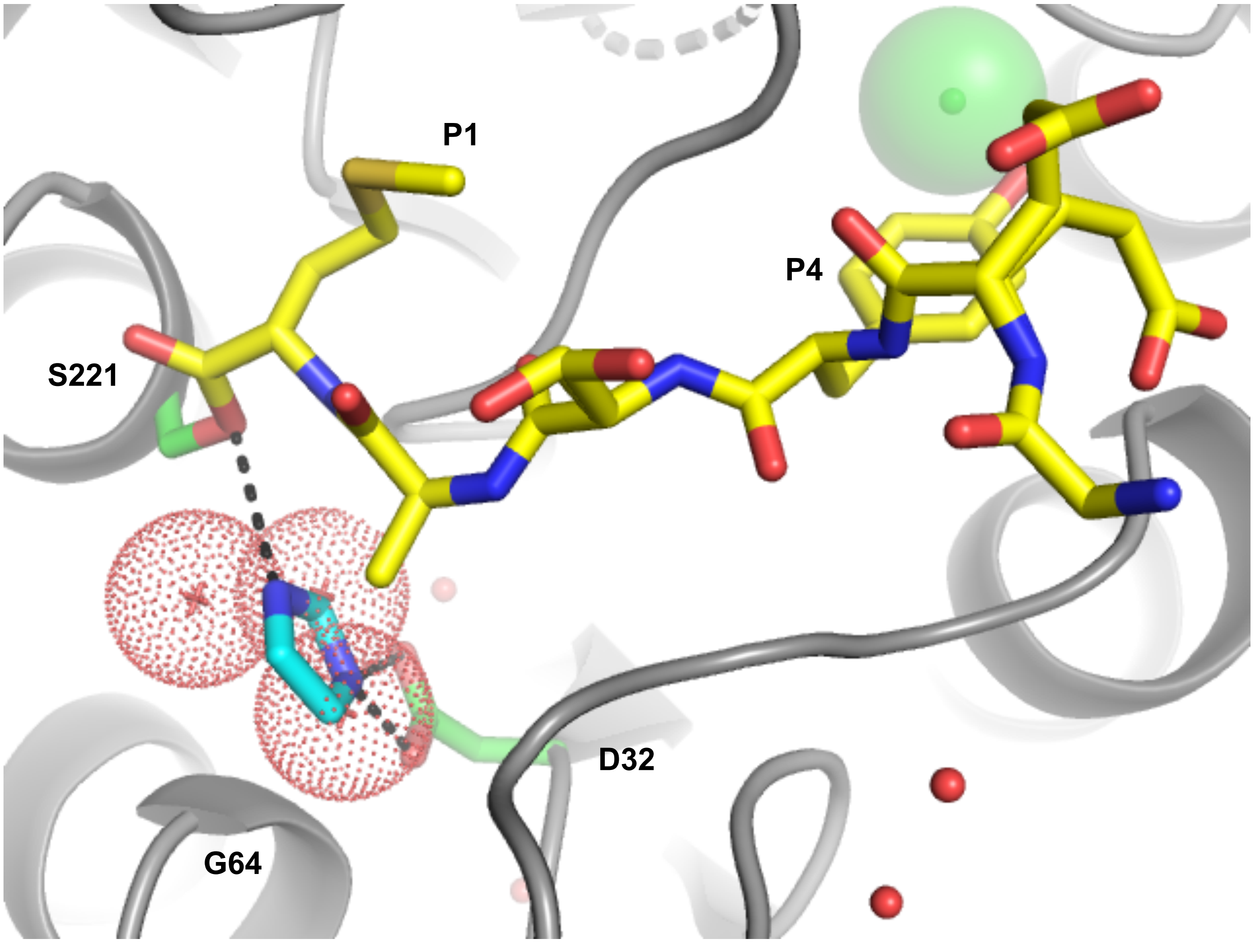


**Fig S7:** *QEEYSAM RASProtease(I) with imidazole in catalytic position*. Structures of RASProtease(I) without imidazole have three conserved waters (dot surfaces) that interact with Oδ1 and Oδ2 of D32, CO of S125, NH of G64, Oγ of S62, and Oγ of S221. When imidazole binds these waters are displaced, the imidazole nitrogens H-bond to Oδ1 and Oδ2 of D32 and Oγ of S221, and the charge relay system is reconstituted.

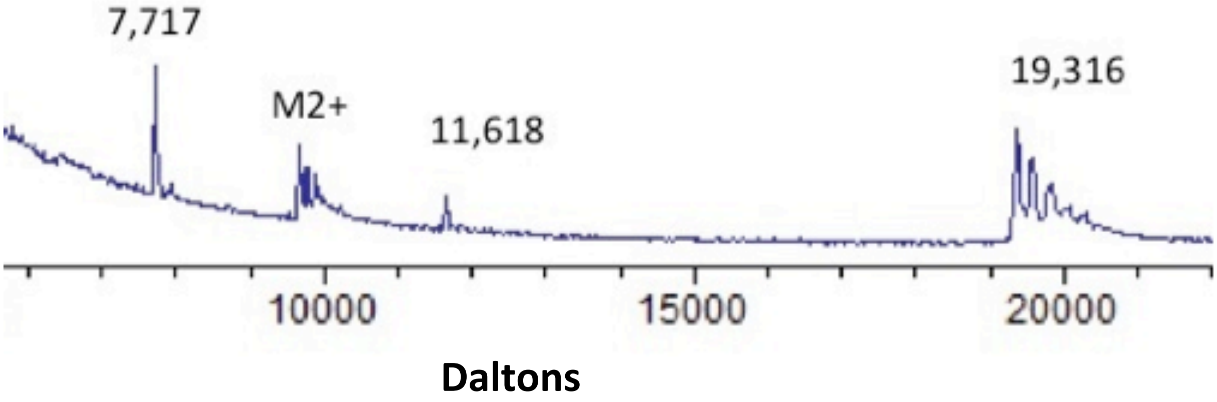


**Fig. S8:** *Analysis of HRAS cleavage.* MALDI analysis (20 hr time point of RAS(GDP) from Fig.5. Intact HRAS is 19,315 Da, the N-terminal fragment is 7,716 Da, and the C-terminal fragment is 11,617 Da.

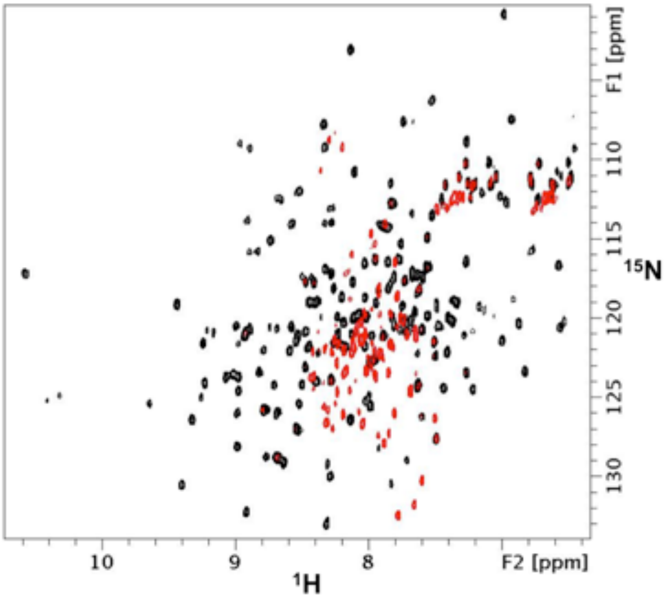


**Fig. S9**: Overlaid 2D $^{1}H$-$^{15}N$ HSQC spectra of HRAS-G12V(GMPPNP) at 100 μM concentration (black) and after treatment with 30 μM protease and 1 mM sodium nitrite at 37°C for 72 h (red). A control spectrum of the HRAS sample with no protease is unchanged under these conditions. The results indicate that cleavage at the YSAM site in the Switch 2 region of HRAS abolishes the globular fold, producing fragments with narrow $^{1}H$ shift dispersion that are consistent with disordered states.

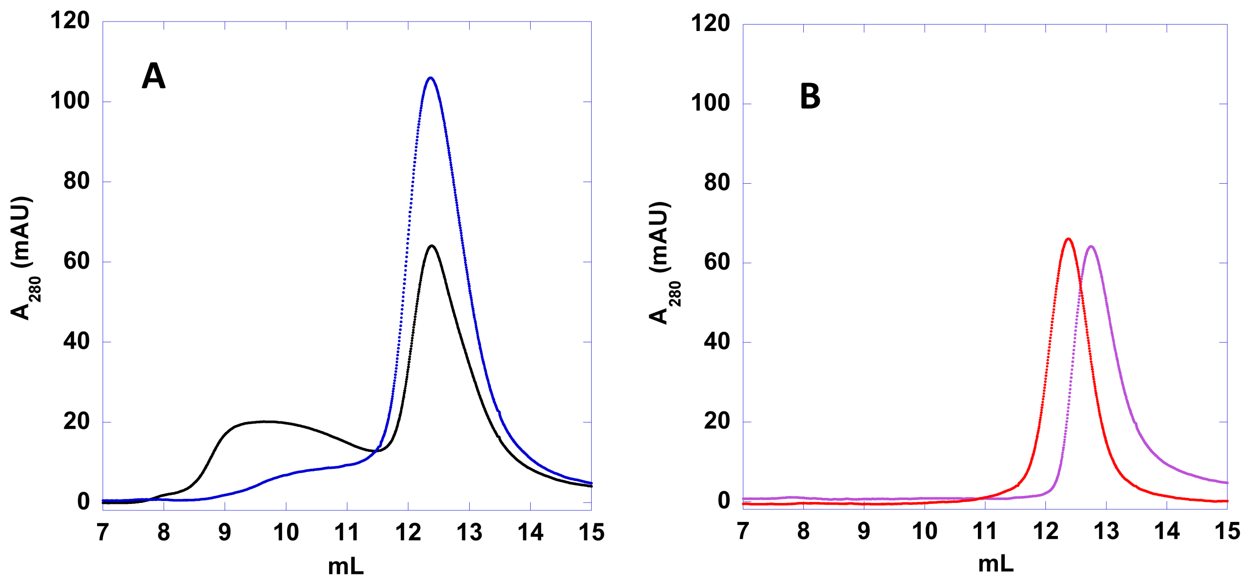


**Fig. S10:** Gel filtration on G75 **A.** Stoichiometric mixtures of RASProtease(I) and RAS. 12.5μM RASProtease(I) - RAS(GDP) is in blue. 12.5μM RASProtease(I) - RAS(GMPPNP) is in black. **B.** Individual proteins. 12.5μM RASProtease(I) alone is in violet. 12.5μM RAS(GMPPNP) is in red. RAS(GDP) elutes identically to RAS(GMPPNP) and is not shown. Based on the fraction bound, $K_S$ (without imidazole) is 6μM for RAS(GMPPNP) and 120μM for RAS(GDP). Molecular weight of RASProtease(I) = 26,438 daltons; RAS = 19,316 daltons (without cofactor).

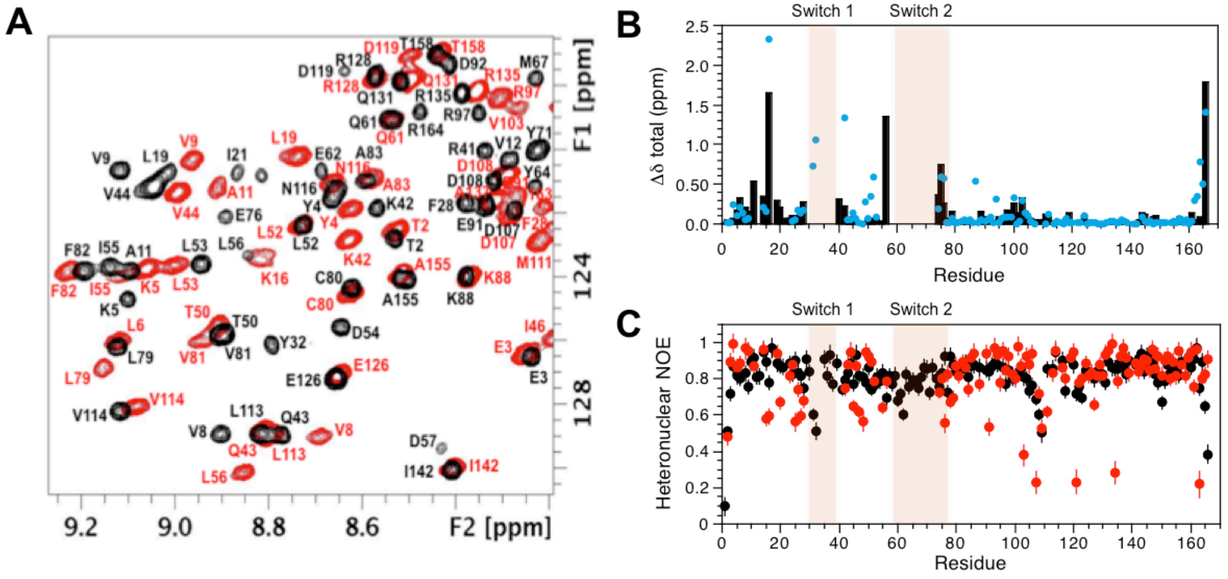


**Fig. S11:** *Structural and dynamic effects of guanosine nucleotide binding on HRAS.* (**A**) Overlaid 2D $^{1}$H-$^{15}$N HSQC spectra showing downfield regions for the G12V variants of HRAS-GDP (black) and HRAS-GMPPNP (red). Backbone amide assignments are shown. F2, $^{1}$H; F1, $^{15}$N. Sample conditions are [HRAS] ~200 μM, 20 mM HEPES, 50 mM NaCl, 5 mM $MgCl_2$, 1 mM TCEP, pH 7.4, 298 K. (**B**) Changes in amide chemical shifts between G12V-HRAS-GDP and G12V-HRAS-GMPPNP (black). Shift perturbations between WT-KRAS-GDP and WT-KRAS-GMPPNP are also shown for comparison (blue) (41, 43). (**C**) {$^{1}$H}-$^{15}$N steady state heteronuclear NOE values for G12V-HRAS-GDP (black) and G12V-HRAS-GMPPNP (red).

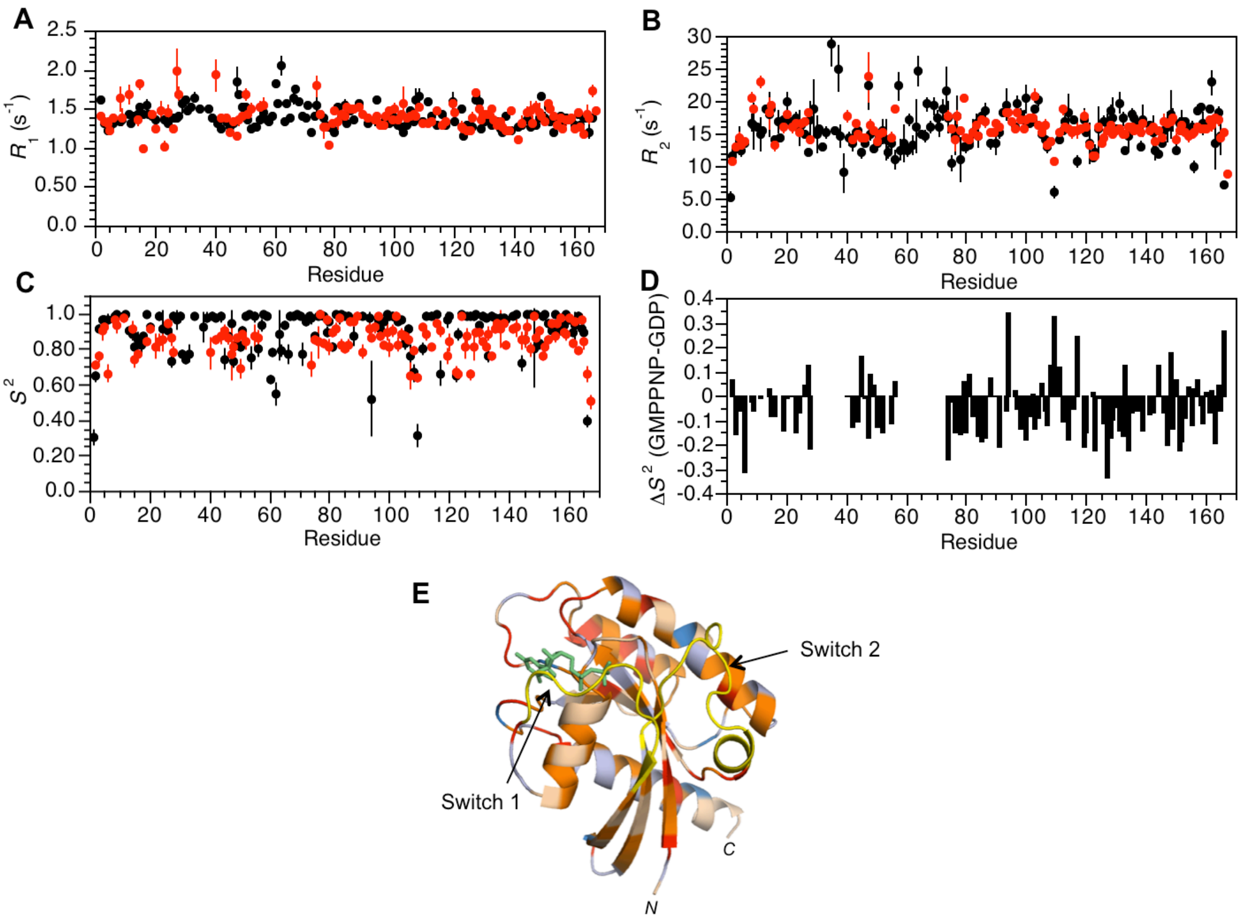


**Fig. S12:** *Changes in HRAS backbone dynamics as a function of nucleotide type.* (**A**) $^{15}$N-Longitudinal ($R_1$) relaxation rates for G12V-HRAS versus residue number. Color-coding for (A-C) is G12V-HRAS-GDP (black) and G12V-HRAS-GMPPNP (red). Error bars indicate ± 1SD. (**B**) $^{15}$N-Transverse ($R_2$) relaxation rates. (**C**) Order parameters ($S^2$) obtained from the $^{15}$N relaxation data, including steady state heteronuclear NOE data in **Fig. S11**, using a model-free formalism. (**D**) Differences in order parameters ($\Delta S^2$ = G12V-HRAS-GMPPNP – G12V-HRAS-GDP) versus residue number. Negative values indicate increased main chain flexibility while positive values indicate decreased flexibility in the GMPPNP-bound form. (**E**) $\Delta S^2$ values mapped onto the structure of HRAS (PDB 2Q21). Color coding is as follows: $0>\Delta S^2>-0.15$, orange; $\Delta S^2<-0.15$, red; $0<\Delta S^2<0.15$, light blue; $\Delta S^2>0.15$, blue. The Switch 1 and Switch 2 regions, which are exchanged broadened in G12V-HRAS-GMPPNP, are highlighted in yellow. The nucleotide is shown in green.

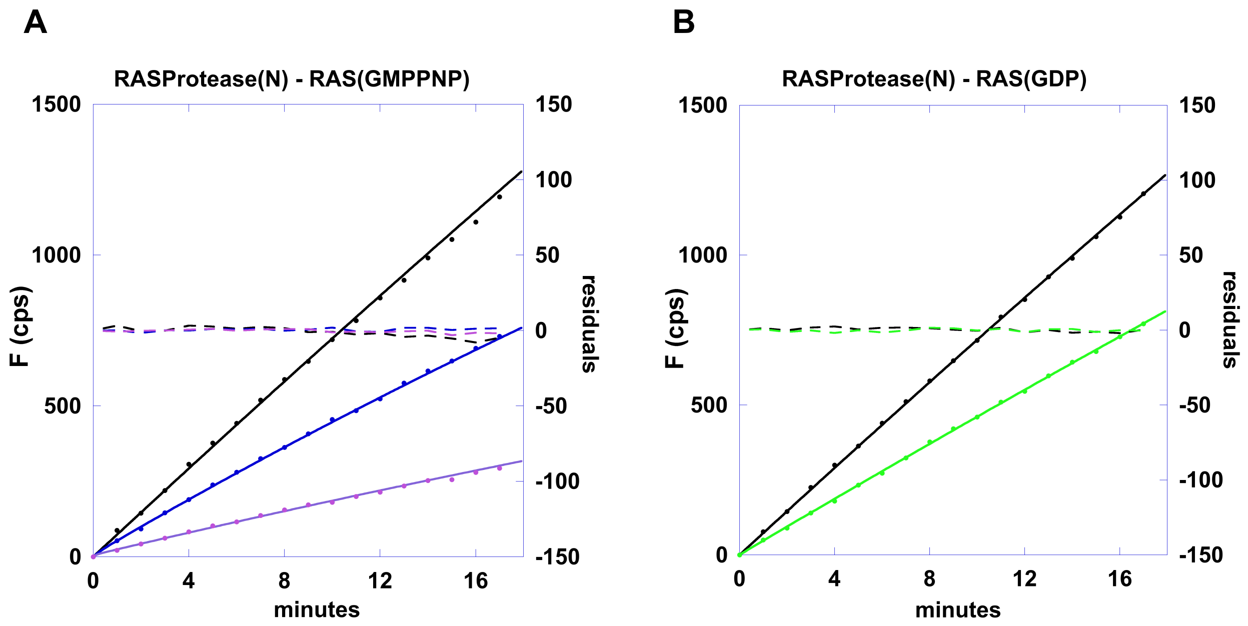


**Fig. S13:** Kinetics of AMC release from QEEYSAM-AMC by 100nM RASProtease(N) in the presence of RAS. **A.** RAS(GMPPNP) **B.** RAS(GDP). The concentration of RAS was 0 µM (black), 1 µM (blue), 5 µM (violet), 10 µM (green) in the presence of 1µM QEEYSAM-AMC and 1mM nitrite. Data points are solid circles. Global fit to mechanism 1 are solid lines. Residuals of data minus fits (5x scale) are dashed lines.

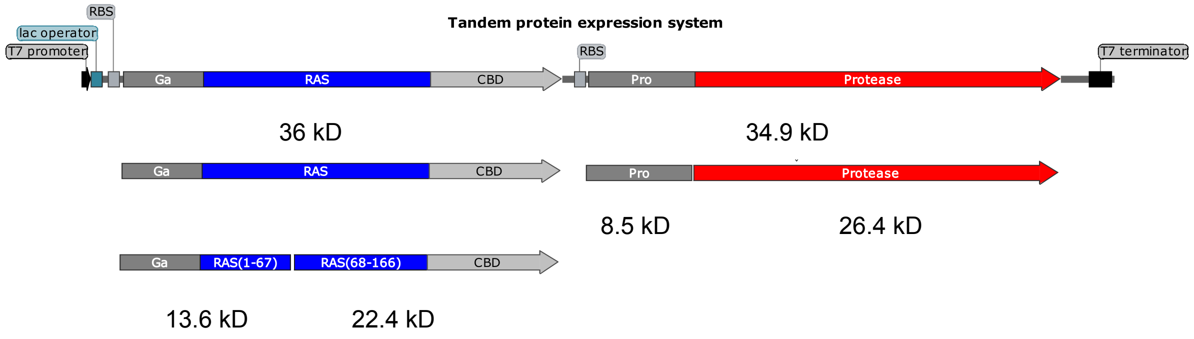


**Fig. S14:** Gene for HRAS fusion protein co-expression with protease zymogen (pro-protease). The intact WT fusion protein is 35,974 daltons. The N- and C- terminal fragments are 13,579 daltons and 22,413 daltons, respectively.

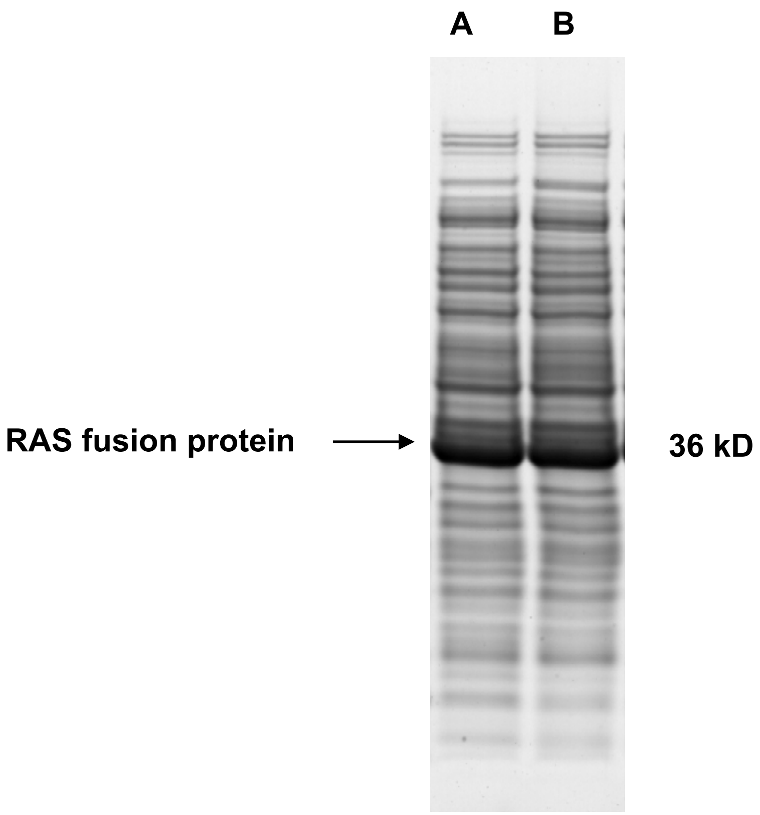


**Fig. S15:** *E. coli* expression of HRAS Lanes A-B: HRAS at 24hrs (**A**) and 42hrs (**B**).

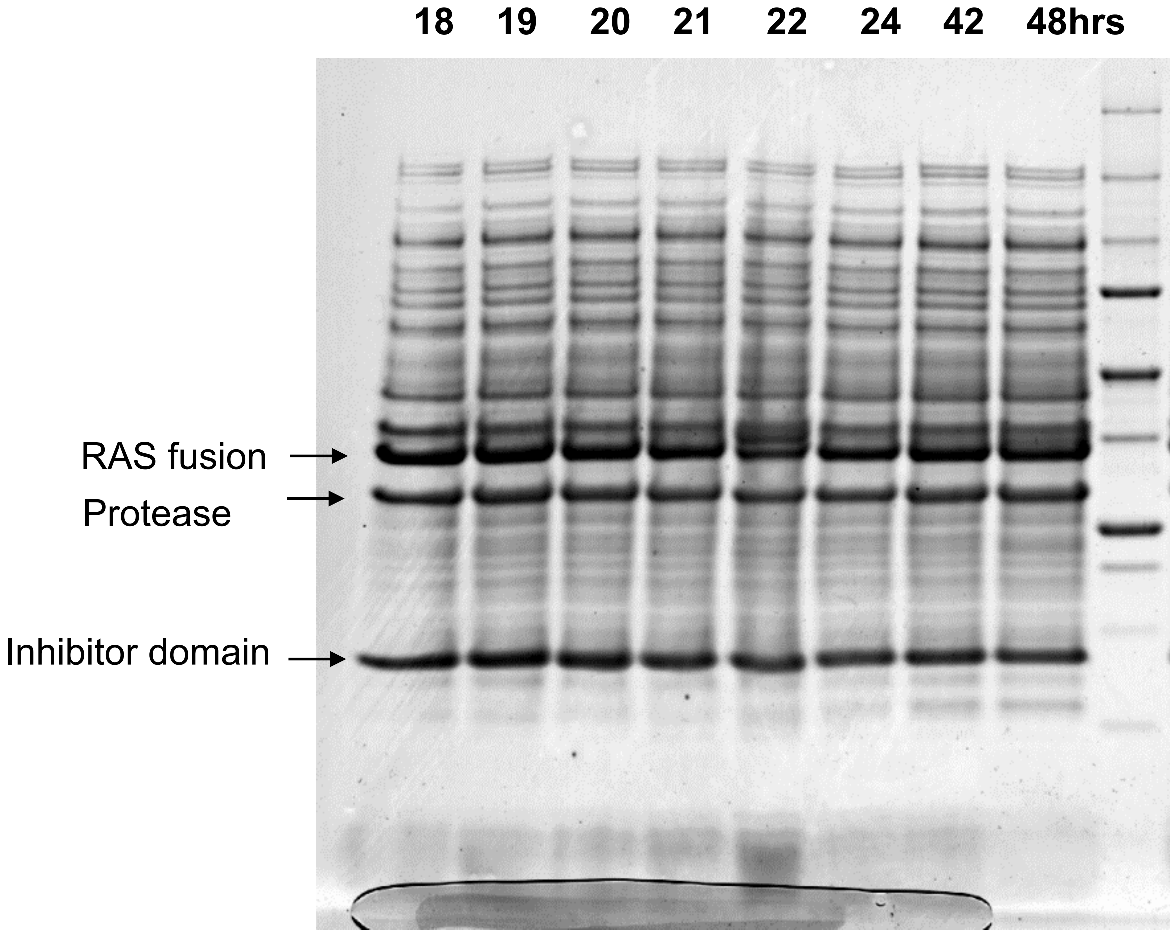


**Fig. S16*: RASProtease(I)*** *No imidazole. Intact RAS fusion protein is 35,974 daltons.* Markers: 250, 150, 100, 75, 50, 37, 25, 20, 15, 10 kDa.

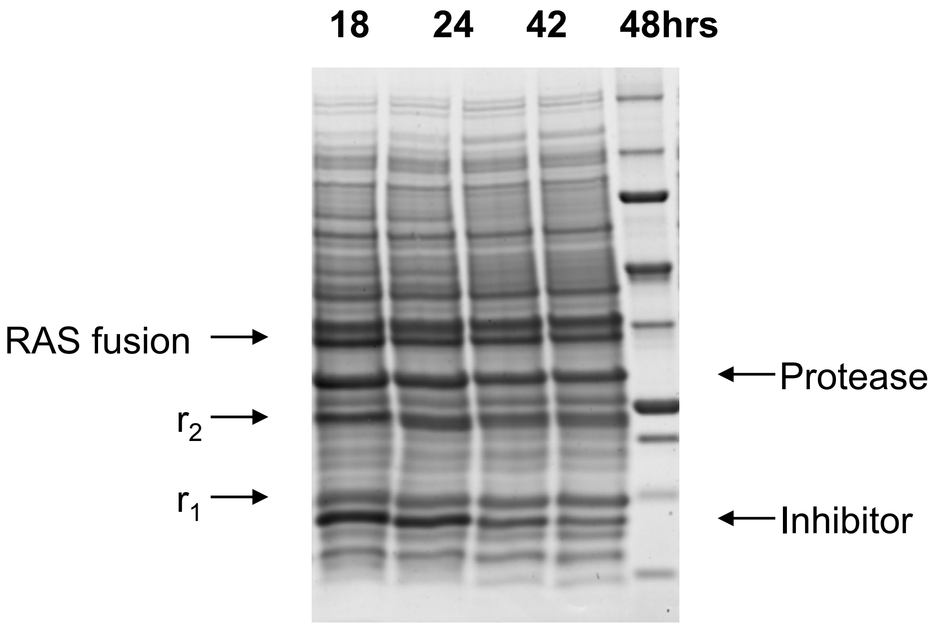


**Fig. S17:** RASProtease(N) zymogen is co-expressed with RAS and endogenous nitrite. Intact RAS fusion protein is 35,974 daltons. The N- and C- terminal fragments are 13,579 daltons (r1) and 22,413 daltons (r2), respectively.

**Supplement: Protein Delivery**

Future evaluation of the therapeutic utility of the RAS-specific protease developed in this study will necessitate delivery within cancer cells. This requires specific recognition of cancer versus non-cancer cells, followed by intracellular transport [1,2]. As such, we started addressing these issues as a feasibility proof-of-concept. With regard to specific uptake by cancer cells, we focused on CD44, a cell adhesion molecule that binds hyaluronic acid and is overexpressed on cancer cells [3,4]. We specifically focused on variants bearing exon 3-encoded ectodomain (CD44v3) given selective advantages: more cancer-specific, longer ectodomain that is more accessible to nanodevices used in drug delivery, and fewer competing interactions with hyaluronic acid [5]. Our pilot results shown in **Figure S18** validate this targeting strategy. For instance, immunostaining with red-fluorescent antibodies showed CD44 overexpression on A431 cancer cells, yet this marker was present on endothelial cells (HUVEC) used as controls, while CD44v3 was cancer specific (**Figure S18A**). Consequently, coating of anti-CD44v3 on the surface of model green-fluorescent nanocarriers (NCs; ≈ 200 nm diameter after coating) resulted in binding on A431 cancer cells at 4°C (a temperature precluding uptake), as revealed by the fact that NCs were accessible to immunostaining with TexasRed-secondary antibody (yellow, green+red dots in **Figure S18B**). At 37°C this method failed to immunodetect cell-associated NCs (green dots) since they were no longer accessible at the cell surface but internalized by cells, a process that was time-dependent and saturable (**Figure S18B**).

Next, we focused on cytosolic delivery of active protease (**Figure S19**). For this, we employed a similarly sized nanodevice built of DNA, called 3DNA®, which we are investigating in parallel through a partnership with Genisphere LLC (Hatfield, PS), and have previously established in other drug delivery settings [6,7]. 3DNA is made from DNA modules, each with two DNA strands that fold into a central dsDNA region, leaving four ssDNA arms [8]. Modules are sequence-engineered for specific arm pairing, so that they self-assemble layer-by-layer into a three-dimensional structure. The outer layer displays ssDNA arms for annealing of oligo-modified cargo and targeting antibodies [8]. As for other DNA nanodevices enabling cytoplasmic delivery, it is believed to low endosomal pH is speculated to change DNA conformation, volume, and amphiphilicity allowing endosome escape of cargo [9]. When A431 cancer cells were incubated at 37°C with targeted DNA-built NCs (unlabeled) and model green-labeled RAS protease (SBT2253) the green protease was visualized spread throughout the cell cytosol, indicating that these NCs favor escape from dot-like endosomes, while incubation with protease alone rendered punctate perinuclear structures (green dots) indicative of uptake and endo-lysosomal retention (**Figure S19A**). Quantification showed increased protease uptake (green sum intensity) and more spread localization (area occupied by green fluorescence) in the presence of these targeted NCs versus incubation of cells with protease alone. These observations were consistent with measurements of activity of protease recovered from cell lysates after introduction of the protease SBT2253 by targeted NCs (**Figure S19B**). This assay allowed us to detect a delivery of ≈ 2 x$10^6$ protease molecules/cell by targeted NCs, without changes in cell viability or morphology.

Finally, we aimed at evaluating whether a RASprotease(N), could cleave its substrate *in situ* within cells (**Figure S20**). Targeted 3DNA NCs were used to first deliver intracellularly the peptide Ac-QEEYSAM-AMC, a specific substrate for this protease, which was modified with amino-methyl coumarin so that its cleavage renders a fluorescent product. RASprotease(N) was delivered the next day into cells containing the substrate, also using targeted NCs. Fluorescence microscopy showed significant detection of fluorescence associated to protease-derived product (**Figure 20A**), which was quantified as increased mean intensity compared to cells which had received protease alone or substrate alone (**Figure 20B**).

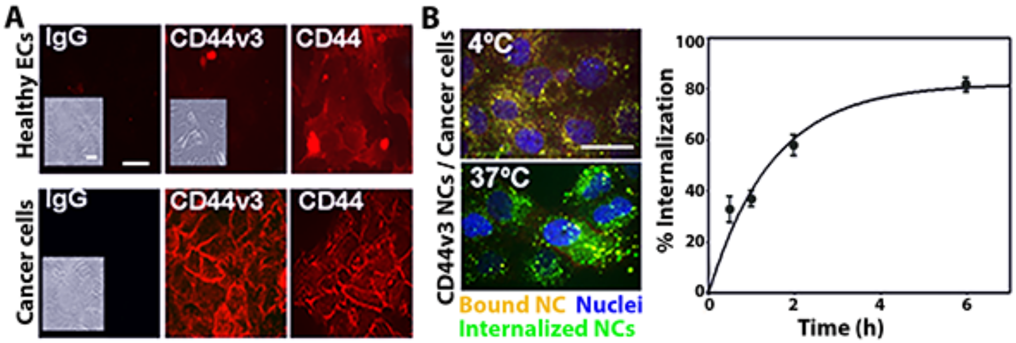


**Figure S18.** Cancer-cell specific targeting and uptake of drug nanocarriers. (A) Expression of CD44 vs. CD44v3 on the surface endothelial HUVEC cells vs. A431 cancer cells, immunostained by 1 h incubation of fixed cells with TexasRed-labeled specific vs. non-specific antibodies. Insets show presence of cells (bright field) for negative samples. (B) Green-fluorescet polystyrene nanocarriers (NCs) coated with anti-CD44v3, incubated at 37°C with A431 cells from 30 min to 6 h, or for 1 h at 4°C as a control. Non-bound NCs were removed by washing, and surface vs. internalized NCs are immunostained using TexasRed secondary antibody, rendering yellow-surface particles vs. green-internalized particles, which were quantified by automatic image analysis as reported (refs). Blue = DAPI-labeled nuclei. Data are average ± standard error of the mean. *p<0.05 by Student's t test. (A,B) Scale bar = 20 µm

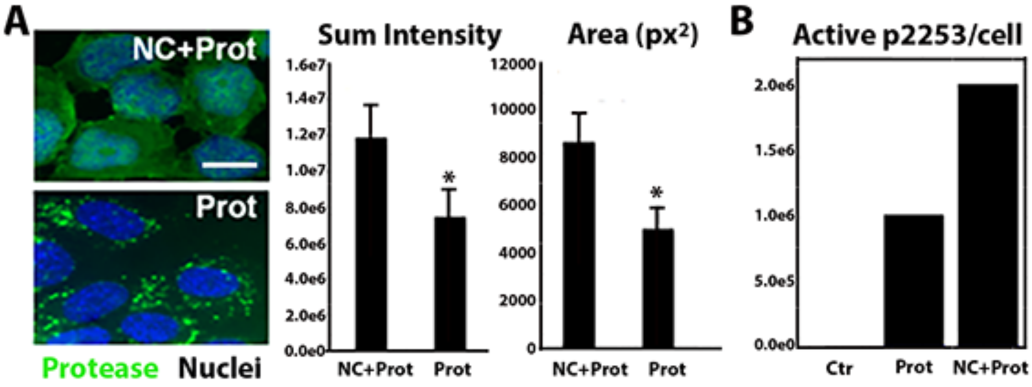


**Figure S19.** Protease delivery to cells. (A) Micrographs and image analysis of A431 cancer cells incubated for 4 h at 37°C with 20 µg/mL green HyLite488-subtilisin protease model (Prot) in the presence vs. absence of 0.4 µg/mL targeted 3DNA nanocarriers (NC). Blue = DAPI-labeled nuclei. Scale bar = 10 µm. Data are average ± standard error of the mean. *p<0.05 by Student's t test. (B) Protease activity was measured *in vitro* in an extract of 500,000 cells (HUVEC) incubated as in (A) with 33 µg/mL green HyLite488-subtilisin protease model (Prot) in the presence vs. absence of 0.66 µg/mL targeted 3DNA NC (control cells are shown; Ctr), from which the mean number of SBT2253 molecules delivered per cell was calculated using a standard curve for SBT2253 activity.

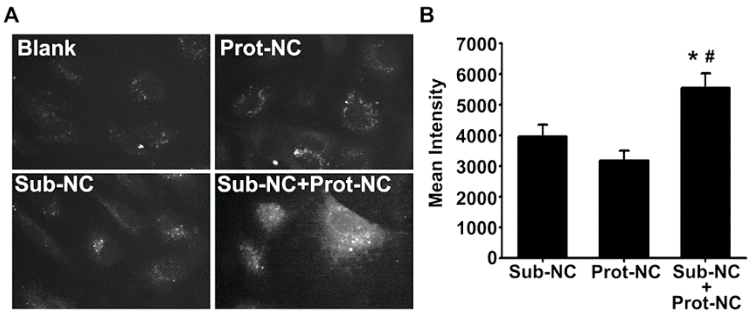


**Figure S20.** Protease activity *in situ* within cells (A) Micrographs of HUVEC cells incubated for 3 h at 37°C with 7.2 µM Ac-QEEYSAM-AMC peptide substrate and 0.4 µg/mL of targeted 3DNA nanocarriers (Sub-NC), followed by washing, cell incubation in control medium for 20 h, and incubation for 1 h with 0.72 µM RASProtease(N) and 0.4 µg/mL of targeted 3DNA NCs (Prot-NC), followed by washing and cell incubation in control medium for 2 h or 23 h. Control involving Sub-NC alone or Prot-NC alone are also shown. Scale bar = 10 µm. (B) Image quantification. Data are average ± standard error of the mean. *p<0.05 by Student's t test.
